# Genetic Basis of Social Structure in the Pastoral Nomads of Central Eurasia

**DOI:** 10.64898/2026.01.27.701587

**Authors:** Ayken Askapuli, Miguel G. Vilar, Maxat Zhabagin, Zhaxylyk Sabitov, Zhaxybay Zhumadilov, Aaron Ragsdale, Theodore G. Schurr, John Hawks, Naruya Saitou

## Abstract

Although the social structure of Central Eurasian pastoral nomads has been described as fluid and *ad hoc* by historians and anthropologists, its potential genetic basis remains poorly understood. To evaluate whether kinship based social organization has biological foundations, we surveyed Kazak populations in Jetisuu, Kazakstan, for genomic diversity with special reference to their kinship structure. We generated genome-wide SNP data (∼750K) using GenoChip microarrays for 80 individuals from four Kazak clans in Jetisuu and 10 Kazaks from other regions. Our results reveal substantial concordance between genetic and genealogical data, with ∼64% of Jetisuu Kazaks sharing the same Y-chromosome haplotype, indicating they have a common paternal ancestry aligning with clan genealogies. By contrast, maternal lineages show remarkable heterogeneity, thereby reflecting female exogamy. Despite this sex-biased admixture pattern, autosomal SNPs reveal no pronounced population structure in Kazaks, suggesting genetic homogenization through ancient admixture followed by continuous gene flow between Kazak populations. Notably, Jetisuu Kazaks exhibit significantly reduced levels of runs of homozygosity (ROH) compared to their sedentary neighbors such as Turkmens, Tajiks, Uyghurs, and Uzbeks. This genomic signature likely results from clan exogamy, which functions as a biological mechanism to prevent inbreeding and maintain genetic diversity. Our findings demonstrate that nomadic social organization represents a sophisticated example of gene-culture co-evolution, where cultural practices have systematically shaped genetic patterns over centuries. These data provide new insights into historical pastoral nomadic societies and offer a nuanced perspective on anthropological debates about kinship authenticity in Central Eurasian nomads.

## Introduction

Central Eurasian nomads have played an important role in the history of Eurasia. They gave birth to well-known historical figures such as Attila the Hun in Europe, Kubilay Khan in China, Ghazan Khan in Iran, and Babur in India^1^. Anthropologists and historians are well familiar with the fact that historical nomadic societies were structured based on kinship and clan identities, which are defined by paternal lineages. For example, the judicial system of the Golden Horde, the northwestern extension of the Mongol Empire, was comprised of four biys (or beys) and the Khan, where the four biys (i.e., judges) represented four major clans of the Golden Horde, and the Khan represents the ruling clan^2^ —*Altan Urug* in Mongol or *Töre,* as Kazaks call it.

In the mid 19^th^ century, the American anthropologist Louis Henry Morgan regarded the human cultural practice of keeping the genealogy of a descent group and organizing people into an intricate kinship-based social structure as one of the earliest examples of human ingenuity after analyzing the social structure of the Native American nations in the League of Iroquois^3^.

The Kazak Khanate emerged as a successor state of the Golden Horde in the 15^th^ century CE, with the ethnonym “*Kazaks*” coming to denote the people comprising the Khanate^4^. Kazaks have a unique type of social unit called the *Jüz*. According to Kazak mythology, all Kazaks trace their ancestry to a single father figure named *Kazak*. His three sons—Akaris, Janaris, and Begaris—gave rise to three *Jüz* (Figure 1).

**Figure 1.**
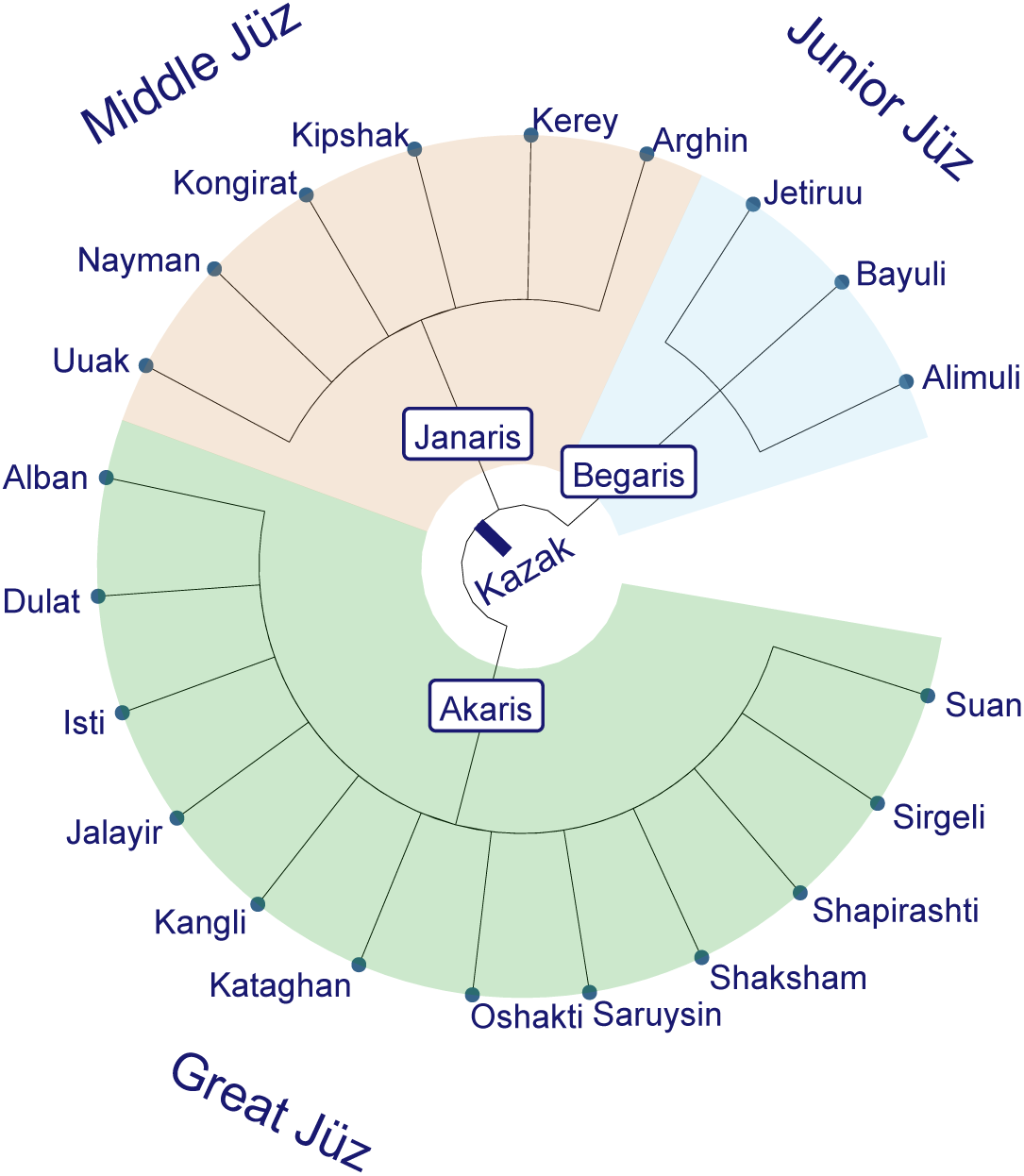
Schematic map of Kazak social structure. The Great Jüz has twelve major clans, while Middle Jüz has six and Junior Jüz has three.

While the existence of clan-based social organization in nomads is well appreciated, the extent to which it is mirrored in the pattern of genetic diversity in these populations has not been adequately evaluated. The historian Peter Golden and other scholars have observed the flexible nature of clan identities within the nomadic societies of Central Eurasia^5^. The ethnic and linguistic identities of these nomadic groups were shaped through various layers of ethno-linguistic influences during the development and decline of the Hunnic, Turkic, and Mongolian empires^5^.

The Mongol Empire, in particular, had a profound influence on the formation of ethnic identities, as Genghis Khan deliberately dismantled nomadic ethnic groups and reorganized them into different military units^5^. Notably, clans such as the Kereys (Kereit), Kipchaks, Jalayirs, and Naymans, which are now part of modern Kazaks and other ethnic groups in Central Eurasia, were identified as separate *nations* during Genghis Khan’s era, as informed by the *Secret History of the Mongols*^6^. According to Rashid al-Din, the medieval Jalayirs were a Turkic nation with ten subbranches^7^.

The nomadic lifestyle granted individuals and groups in steppe societies the freedom to relocate, allowing them to easily change political alliances. This mobility contributed to a dynamic social structure without strong ties to specific territories. As a result, kinship and genealogical frameworks developed from an *ad hoc* need for political alliances^8–10^. Some anthropologists believe that the organization of a people into unilineal descent group was a product of state administration in the most regions of Inner Asia, asserting that the clan identity in Mongolia—*obeg*—was imposed on Mongols by the Manchu Empire for administrative convenience^11^, despite the fact that the Mongolian genealogy and clan identity already existed at the time of Genghis Khan^6^.

In this context, Chaix and colleagues analyzed the Y-chromosome short tandem repeat (STR) profiles in Turkic-speaking populations in Karakalpakstan, a republic within Uzbekistan, including Kazaks, Turkmens, Uzbeks, and two Karakalpak clans^12^. They did not find any significant correlation between the patterns of paternal genetic diversity and clan organization, leading them to conclude that the genealogical records of these groups may be fictitious^12^. However, the Kazak participants of the study were not identified by clan but instead grouped into a single Kazak population.

By contrast, other genetic research on Kazak populations showed that most members of the same clan shared the same Y-chromosome haplogroup^13,14^, suggesting a genetic underpinning for clan identity among Kazaks. However, these previous studies reported only Y-chromosome haplogroup frequencies and failed to examine possible connections between paternal lineages and the social structure with reference to genomic profiles. In addition, the lack of high-resolution phylogenetic analysis of Y-chromosomes raised questions about making definite association between genetics and genealogy, as most of the studies only analyzed a limited number of STRs and binary SNP markers.

In this study, we investigated whether the social structure of nomadic peoples has any genetic basis through the genetic analysis of four Kazak clans from the Jetisuu region. This region is well-suited for pastoral nomadism as multiple rivers and streams arise in the Tängirtau Mountains (Tianshan) and flow into the Lake Balkash. Among these watercourses, the largest is the Ile River. The mountain slopes in the south provide summer pastures, while the lowland areas near the Lake Balkash are hospitable for herding during the winter times. Archaeological studies indicate that the nomadic subsistence tradition of the region can be traced back to the Bronze Age^15^.

The Great *Jüz* has twelve clans in total (Figure. 1). Seven of them (Alban, Dulat, Isti, Oshakti, Sarüysin, Shapirashti and Suan) are grouped together as Üysin. We surveyed four clans from the *Great Jüz,* including Jalayir, Shapirashti, Alban, and Suan, which mainly live in the Jetisuu region. Some historians believe that the Üysin trace their ancestry to Wusun, an Iron Age people who inhabited the Ile River basin, i.e., Jetisuu, with the Wusun believed to be one of the ancestral sources of Kazaks^16^. The head of the state of Wusun was called [kunmo] or [kunmi], where *kun* could be the Turkic word *kün*, meaning *the sun*^16^. However, a direction connection between the Iron Age Wusun and Üysin is highly unlikely (Peter Golden and Christopher Atwood, personal communications, 2025). Some historians connect the ethnonym Wusun with Old Indic *asvin*, the appellation of twin equestrian gods, suggesting an Indo-European origin for the Iron Age Wusun people. They draw on the fact that the Chinese character *Wu* was pronounced as [a] in ancient Chinese, despite it now reads [wu]; in addition, the Wusun people had Caucasian appearance as informed by Chinese annals^17^. Yet, other historians associate the Üysin of modern Kazaks with the medieval Mongol people Üshin^18^(also spelled *Hüshin)*.

As for the origin of Jalayirs, another clan in the Great *Jüz* that does not belong to Üysin, historians believe that they are the descendants of medieval Jalayirs who played important roles in the Mongol Empire. For instance, Mukhali the Jalayir was a general of Genghis Khan who led the eastern front in China^1^. He was from the Jait clan of Jalayirs^7^; and in the west, many Jalayirs served in the Il-khanate in Persia and the Middle East^19^.

In the context of pastoral nomads in Central Eurasia, we define *clan* as a patrilineal kin group that is patriarchal, patrilocal, and exogamous. Patrilocality is a feature shared by most human societies^20^. If clans are formed as an *ad hoc* alliance between different nomadic groups, such clans should harbor heterogeneous paternal lineages, thus ruling out the possibility of shared paternal ancestry among its members. By contrast, if a clan represents a genetically related patrilineal descent group, then most male members of the clan should share the same paternal lineage, indicative of descent from a common paternal ancestor. However, other paternal lineages may also exist in the same clan due to admixture and other demographic events.

Ethnic groups in Central Eurasia can largely be categorized into two groups—pastoral nomads and sedentary agriculturalists. The two groups differ in subsistence patterns as well as cultural practices, although the boundaries between the two are far from clear-cut.

A research group from China^21^ carried out a survey of marriage patterns in several ethnic groups, including nomadic Kazaks and sedentary Uyghurs, in the Ile region in Xinjiang, which is situated in the upper reaches of the Ile River, a natural extension of the Jetisuu across the Kazak-Chinese border. The survey revealed that the highest rates of consanguineous marriages occurred in Dungans and Uyghurs (8.1% and 8.23%, respectively), while the Sibe had a moderate rate of 4.66%; Kazaks a low rate of 2.87%, and Mongols had the lowest (0.45%). Here, the former two groups are sedentary agriculturalists while the latter two are pastoral nomads (Figure 2H). Dungans and Uyghurs, the groups with the highest rates of consanguinity, live in the same region and have similar subsistence patterns. Despite these facts, they are linguistically distinct from each other, with Uyghurs speaking a Turkic language, and Dungans speaking Chinese, and also genetically differentiated, with Dungans having greater genetic affinity with East Asians (e.g., Han Chinese)^22^. However, a unifying feature of the two was their shared belief in Islam. By contrast, their Muslim but traditionally nomadic neighbors, Kazaks and Kirghiz, tend to have more superficial religious beliefs characterized by syncretism, blending Islam with traditional beliefs, rather than following scriptural institutions^23,24^. In the 19^th^ century, the Russian Empire implemented a systematic policy aimed at converting nomadic Kazaks into “true” Muslims^25^.

**Figure 2.**
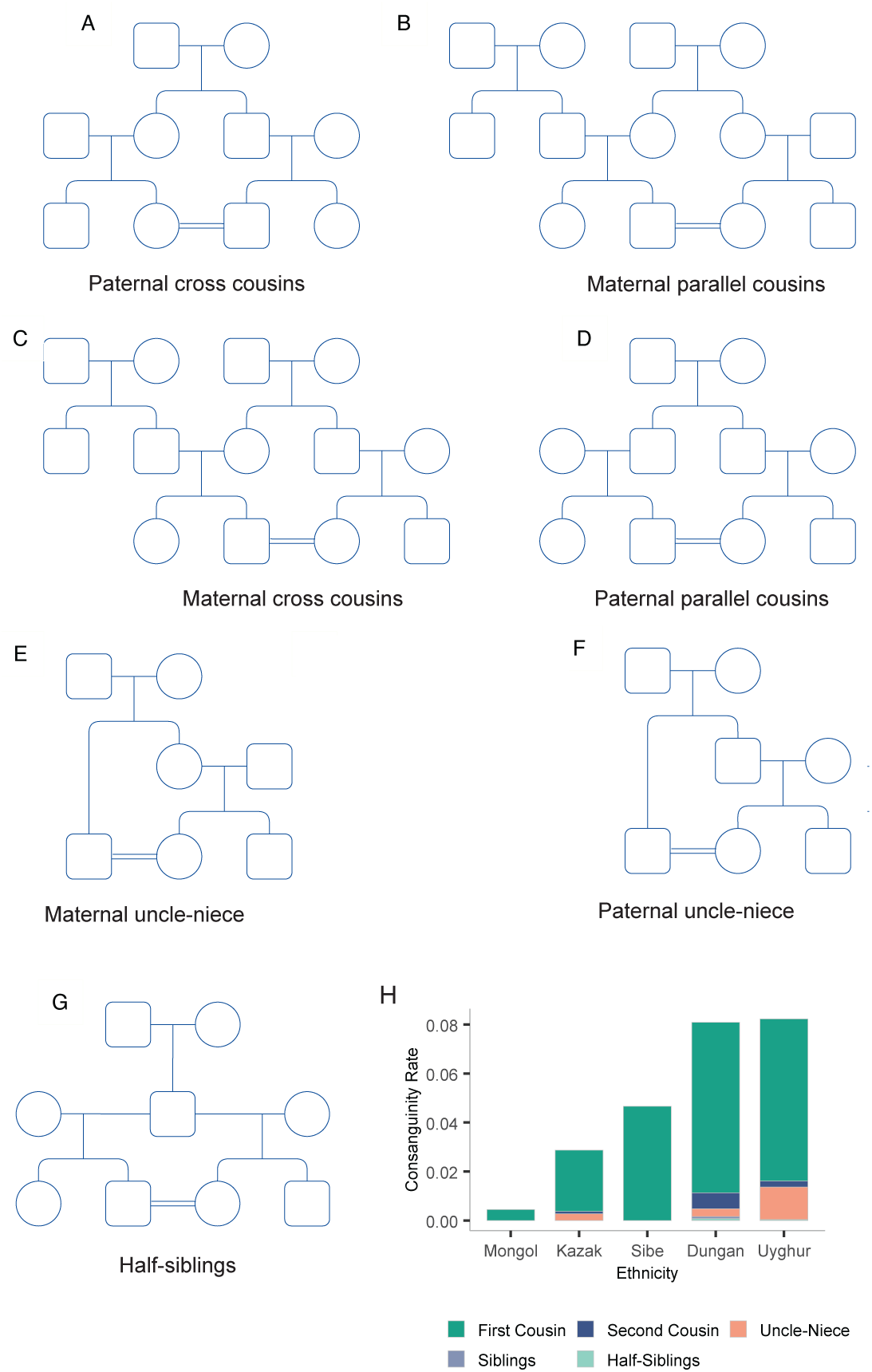
Consanguineous marriage patterns and rates found in Central Asia. The bar plot was made based on the survey data from Ai and colleagues (1985)^21^.

In Kazaks and other pastoral nomads in Central Eurasia, knowledge of kinship structure and genealogy serves as guidance for marriage. Kazaks practice exogamy with a cut-off of seven-generation patrilineal relatedness, while for the maternal relatedness no such strict rule is applied.

Interestingly, consanguineous marriage patterns vary among nomadic and sedentary groups as well (Figure 2). The most frequent kind was first cousin marriages in all ethnic groups surveyed. While the frequency of paternal parallel first cousin marriages (Figure 2D) was 18.09 % in Uyghurs and 4% in Dungans, no such cases were observed in Kazaks, Mongols, and the Sibe. Paternal uncle-niece marriages (Figure 2F) also occurred in Uyghurs (5.71%) and Dungans (2%) but not in Kazaks, Mongols, and the Sibe.

The consanguineous marriage patterns seen in Uyghurs and Dungans are common in the Muslim communities of West Asia and Middle East. In Turkey, consanguineous marriages account for 22% of all marriages, with the paternal parallel first cousin marriage being the most common type^26^. In Iran, the overall rate of consanguineous marriage is 38.6%, with the highest rate being found in the Baloch (59.9%), among whom paternal parallel first cousin marriage was the most common type^27^. In Arabic communities, the most common type of consanguineous marriage is the paternal parallel first cousin marriages as well, which account for 35-65% of consanguineous unions^28^. In the Arabic culture, only the “milk siblings”, those born to the same mother, are not allowed to marry^29^.

Even though consanguineous marriages occur at high frequencies in Muslim communities of Islamic countries in Middle East, North Africa, and West & South Asia^28,30^, such close kin marriages also occurred at elevated frequencies in 19^th^ century Europe^31^. A well-known example is the consanguinity in the family of Charles Darwin^32^. It is thought that consanguineous marriages in Europe began to disappear due to industrialization^33^ and the decrease in fertility rates^28^.

First-cousin marriage is also allowed in Buddhism^34^. The nationwide rate of consanguineous marriages in Japan was reported to be 3.88%, with lowest rate (0.03%) in the north and highest in the south (7.89%) of the country^35^. Also, Hindu believers in South India have a consanguineous marriage rate of 30%, of which over 20% being uncle-niece unions^36^.

Motivated by these kinds of issues, we evaluated the impact of social structure and marriage customs of nomads on their genetic diversity and compared them with the features of sedentary agriculturalists. More specifically, we generated and analyzed genomic SNP data of 90 Kazak individuals, including 80 individuals from the four clans of the Great *Jüz* and 10 Kazak individuals from other regions of Kazakstan. In what follows, we describe how nomads organized themselves and explain the outcomes of social organization and customs as reflected in their genomes.

## Subjects and Methods

### Fieldwork and sample processing

Fieldwork was performed in Jetisuu (Figure 3) under the framework of a project established at Nazarbayev University (NU) in Astana, Kazakstan, with a research ethics review and approval by the Ethics Committee at the Center for Life Sciences (CLS), National Laboratory Astana (NLA), NU. We have undertaken several fieldwork sessions in Jetisuu (literally meaning *seven rivers*), as described previously^37^.

**Figure 3.**
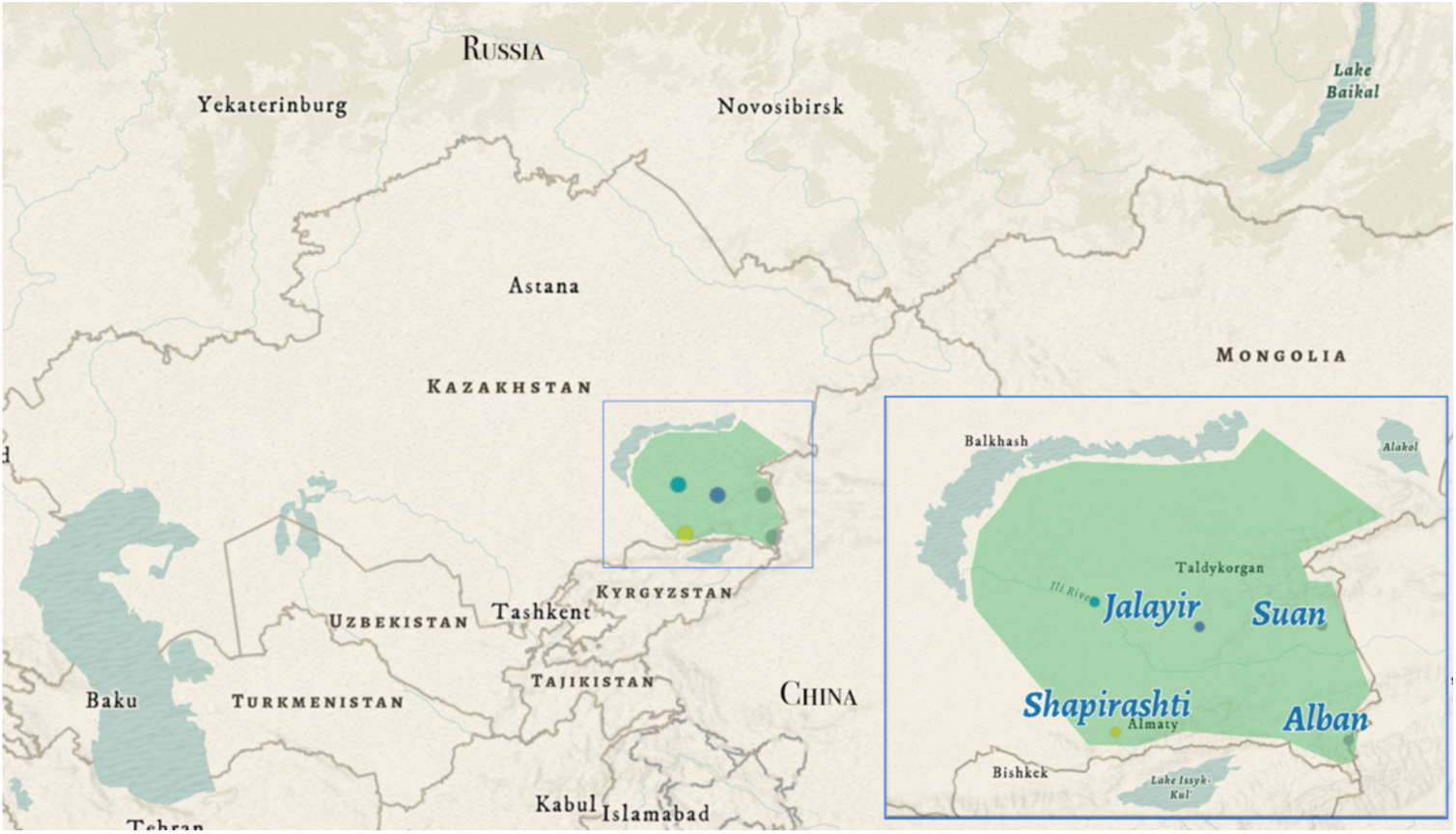
Map of Jetisuu and the Sampling Sites. The green area in the blue square depicts the Jetisuu region within the border of Kazakstan. The colored dots represent approximate location of the sampling sites.

All participants were enrolled in the project by signing a written consent form, and thereafter, completing a genealogical questionnaire. Blood samples were collected via vacutainers (BD), with help of medical nurses. DNA was extracted from blood using a salt extraction method^38^, with modifications or a commercial kit (e.g., Qiagen); blood and DNA samples were stored at - 86℃ at CLS, NU as described previously^37^.

### Microarray genotyping and data compilation

#### Genotyping

Ninety Kazaks, including 80 individuals from Jetisuu and 10 individuals from other regions of Kazakstan, were genotyped using the GenoChip 2.0 microarray (Illumina iSelect HD)^39^ under the framework of the Genographic Project. This GenoChip called ∼750,000 SNPs throughout the genome for each individual, including markers for the mitochondrial DNA (mtDNA), autosomes (Chr1-22), and the X- and Y-chromosomes.

#### Y-chromosome data analysis

This microarray data had ∼14,000 SNPs from Y-chromosomes, of which∼6,000 intersected with the set available in the 1000 Genomes Project (KGP) Y-chromosome dataset^40^. After removing either monomorphic or tri-allelic variants and excluding sites with more than 10% missingness in the Jetisuu dataset, we were able to merge it with the KGP Y-chromosome data^40^. The final Y-chromosome dataset contained 1323 individuals and ∼1,100 SNPs per individual.

#### Autosomal DNA analysis

The autosomal data (Chr 1-22) from the Jetisuu individuals was subjected to QC via PLINK^41^, with the parameters [--geno 0.1 –maf 0.05]. As a result, the analysis of DNAs from 88 Kazak individuals, including 78 from Jetisuu and 10 from other regions, yielded 558,959 SNPs after cleaning the dataset. The two individuals (SH002 and SH004) were removed from the dataset, since they were outliers based on both the non-recombining portion of the Y-chromosome (NRY) and autosomal data. The refined Jetisuu dataset was merged with a dataset (N=925) from Lazaridis and colleagues^42^ to produce a merged dataset containing 1,013 individuals (from ∼70 populations) and 114,376 SNPs. We created a separate LD (linkage disequilibrium) pruned dataset via PLINK v1.9^41^ using the parameters [-- idep-pairwise 50 5 0.2]. The LD pruned dataset, containing 1,013 individuals and 59,102 variants, was used to conduct PCA analysis in PLINK v2^43^ and admixture analysis via Neural Admixture^44^. Sample information is given in Table S1 and S2.

### Y-chromosome phylogeny

Y-chromosome haplogroups were called via yHaplo (v2.1.14)^45^ using a dataset of 1323 Y-chromosomes in VCF format as an input file. The dataset was compiled by merging our dataset (N=90) with the KGP Y-chromosome data (N=1233)^40^. Manual inspection of diagnostic NRY haplogroup markers enabled us to further identify detailed haplotypes of some Jetisuu samples. The phylogenetic trees were reconstructed with RAxML-NG^46^ using the final dataset in FASTA format, which was generated by concatenating SNPs to form pseudo-DNA-sequences via VCF-Kit^47^. The final tree (Figure 4A) was visualized with the R package ggtree^48^, with some modifications, i.e., trimming some branches (e.g., O and R) and adjusting branching patterns of some clades (e.g., A and B), based on a previously published Y-chromosome tree^40^.

**Figure 4.**
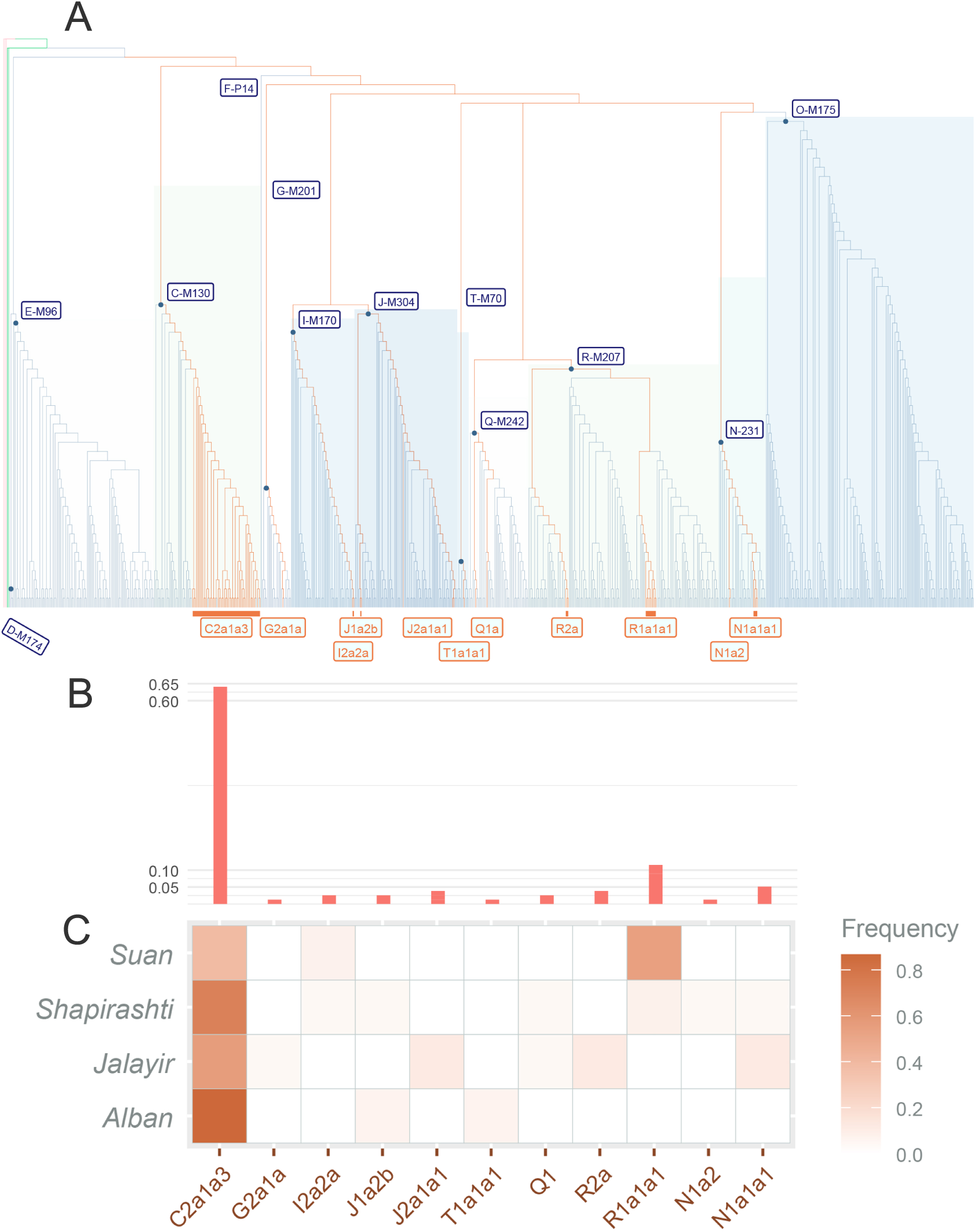
Y-chromosome phylogenetic tree and the frequencies of haplogroups found in Jetisuu populations. **Note: A.** A Y-chromosome phylogenetic tree, where orange branches represent paternal lineages found in the Jetisuu Kazaks while blue branches are Y-chromosome lineages from the 1000 Genomes Project; additionally, the blue branches include 10 Kazak individuals from this study; **B.** Frequencies of lineages found in the Jetisuu Kazaks, with the positions of the bars corresponding to the tree branches above and the tabular heatmap columns below. **C.** The tabular heatmap shows the frequencies of Y-chromosome lineages within each of the four clans. Here J2a1a1 includes three individuals. One of them had higher missingness in genotype and yHaplo called it as basal J. Our manual inspection has confirmed that it belongs to J2a; therefore, we tentatively grouped this individual with other two, since all from the Jalayir.

### Principal component analysis (PCA)

PCA was carried out using PLINK v2.0^43^ with the LD-pruned dataset in PLINK binary format as an input. PLINK calculated 10 principal components and, for plotting purposes, we used the first two, since they explained the most genetic variation (68.3%). PCA results were graphically represented using the ggplot2 package of R^49,50^.

### Admixture analysis

The LD-pruned dataset in PLINK binary format was used for admixture analysis via the Neural Admixture program (v1.3.0)^44^. We used the neural-admixture *train* model with the parameters [--min_k 3 --max_k 9 –initialization pckmeans –max_epochs 5]. This program is much faster than the commonly used ADMIXTURE program^51^, although the current version of Neural Admixture does not have the functionality to cross-validate K values. The admixture clusters were visualized with the R package pophelper^52^.

### Analysis of runs of homozygosity (ROH)

We searched for runs of homozygosity (ROH) using PLINK v1.9^41^. With the –homozyg functionality, using default parameters, we identified ROH segments equal to or longer than 1 Mb that had at least 100 SNPs. The dataset with 1,013 individuals and 114,376 SNPs in PLINK binary format was used as an input. The ROH results were visualized with the ggplot2 package of R^49,50^. The results of this ROH analysis are given in Table S2.

### Identity by descent (IBD) analysis

We used the program Identical by Descent via Identical by State (IBIS)^53^ to detect IBD segments between individuals and HBD (Homozygosity by Descent, i.e., ROH) segments within individuals. IBD segments of 7 cM or longer with at least 436 SNPs were called with the parameters [-ibd2 -hbd -mL 7 -mt 436 -er 0.004]. The input file was the same dataset used for ROH analysis via PLINK. IBIS rapidly detects IBD and HBD segments in unphased genomic data^53^. The results of IBIS were visualized with the ggplot2 package of R^49,50^. The information on the IBD segments and the inbreeding coefficient is included in Table S3.

### Kinship

Kinship among the Jetisuu individuals was evaluated by creating a distance matrix based on the kinship coefficients among individuals. Kinship coefficients (Table S4) were calculated by the *IBIS* program^53^ based shared IBD segments. A kinship tree (Figure S5) was built via the Neighbor-Joining method^54^ , implemented in the R package *ape* (v5.8.1)^55^. The kinship tree was visualized with the R package *ggtree* (3.12.0)^48^.

### Statistical tests

For testing the significance of the differences in ROH levels, we performed Mann-Whitney U Test, which is also known as Wilcoxon Test, using the wilcox.test function in R^50^, with p-value < 0.05 as a cutoff for statistical significance. In addition, we used Fishers Exact Test for comparing the proportion of individuals with long ROH. Fishers Exact Test was carried out via fisher.test in R^50^.

## Results

### A single Y-chromosome lineage characterizes the Jetisuu Kazaks

Our analysis of the paternal genetic profiles of the Jetisuu Kazaks indicated that their paternal lineages were relatively homogenous. Just a single haplotype, C2a1a3-F1918, i.e., the C3 Star Cluster (C3*), accounted for ∼64% of the paternal lineages observed (Figure 4). Among them, however, three samples (AL006, AL011, and SH025) did not have a derived allele for the marker F1918, which could be due to a genotypic error. This finding contrasted sharply with the elevated levels of maternal genetic diversity observed in these populations, which had diverse sets of East Eurasian (A, B, C, D, F, G, and Z) and West Eurasian (H, HV, V, W, I, J, R, and X) lineages. In fact, the highest frequency of a maternal lineage was just 16% (D4), while the rest occurred at a frequency of 7.5% or less^37^.

Among the minor paternal lineages found in the Jetisuu Kazaks, only the frequency of R1a-M17 (i.e., R1a1a1) exceeded 10%, with others being present at 5% or less. Interestingly, the minor haplogroups were clan specific. For instance, R1a1a1 individuals were mostly found in Suans, with an additional two individuals from Shapirashti having this lineage. R2a was found only in Shapirashti, and T1a1a1 only in Albans (Figure 4C).

However, our data may not fully reveal the pattern of paternal diversity in these clans because of the small sample sizes. As an example, a greater number of haplogroups were present in Jalayir and Shapirashti compared to the other two clans, most likely due to sample size differences (∼25 vs. 13 and 15). The frequency of R1a-M17 was slightly higher than that of C2a1a3-F1918 in Suans, a result that was consistent with a previous genetic study of Suans^56^. The two haplotypes also occur together in Kirghiz^57,58^, who have higher frequencies of R1a-M17 compared to C2a1a3-F1918.

While our results reveal C2a1a3-F1918 as the major paternal lineage of Jetisuu Kazaks, it is necessary to analyze a large set of samples for characterizing minor haplogroups in these populations. The results of current study support the Mongolian Üshin origin for the Üysin union of the Great Jüz, since C2a1a3-F1918 (i.e., C3*), the lineage associated with Mongol expansion^13,59^, characterizes all four clans analyzed. Keeping these findings in mind, we will now describe each of the paternal lineages found in the Jetisuu Kazaks.

### C2a-F1918

C2a-F1918 (i.e., C2a1a3), a major branch of C3*, was previously hypothesized to be associated with Genghis Khan. It has been found in populations residing across a wide geographic area spanning from the Caspian Sea in the west to the Pacific coast in the east^59^. While C2a1a3 occurs at high frequencies in Kazaks^13^ and Hazaras^59^, it is also found in various Mongolic speaking groups at moderate frequencies^60^, while occurring at lower frequencies in various other ethnic groups across Eurasia^59^. C3* is also one of the founding paternal lineages of Native Americans^61^, with C2a2b-MPB373 occurring in native populations in South America^62^. Besides the Jetisuu Kazaks, C2a1a3-F1918 occurs at high frequencies in the Kerey and some other clans of Kazaks^13,63^. In Jetisuu, C2a1a3-F1918 occurs at high frequencies in all four clans of the Great Jüz (Figure 4).

### R1a-M17

Haplogroup R is found commonly in Europeans, with a decreasing clinal distribution from the west to the east, although its oldest branches are found in Central Asia^64,65^. R1 is among the most thoroughly investigated of all human paternal lineages. R1a and its sister lineage, R1b, are seen in all Indo-European-speaking populations and occur at high frequencies in Europe, India, and Central Eurasia, including southern Siberia^64–67^. The spread of haplogroup R1b is believed to be associated with the Kurgan Culture people, also known as the Yamnaya, who spread Indo-European languages^67^. However, Yamnaya individuals have been found to have only R1b haplotypes, whereas earlier Eastern European Hunter-Gatherers (EEHG) carried both R1a and R1b Y-chromsomes^67^. The earliest record of R1a appears in Mesolithic burials of Europe belonging to the EEHG^68^. In Central and West Asia, R1a is relatively frequent, with its highest diversity being found in Iran, indicating its possible geographic origin there^64^. However, the Bronze Age populations of Iran did not have R1a and R1b^66^. The highest frequencies of R1a-M17 occur in Altaians (∼12–50%)^69^ and Kirghiz (∼45–63%)^58,65^, both speaking the Kipchak branch of Turkic languages, while also appearing in the Persian speaking Tajiks (19–64%)^65^. In Kazaks, R1a-M17 (i.e., R1a1a) occurs at relatively low frequencies, with the highest frequency occurring in the Suan clan (31.7%)^56^. In our dataset, seven out of 13 Suan individuals (i.e., 53.8%) harboring this haplogroup.

### R2a-M124

The global frequency of R2a-M124 is much lower than those of R1a and R1b. This haplogroup is found in Central Asia, Caucasus, Middle East, and South Asia^65^. In Central Eurasia, it occurs at low frequencies (2-9%) in various ethnic groups, with some exceptions (e.g., 53% in Sinte Romani population in Uzbekistan)^65^. Among Kazaks, it occurred at its highest frequency (17%) in the Töre clan^70^.

### N1a-M178

Haplogroup N, the sister clade of haplogroup O, likely originated in southeast Asia^71^. N1a-M178 (N1a1a), the most frequent subbranch of N-M46 (aka N-TAT) in Siberia, is the signature haplotype of Sakha (Yakut), a Turkic speaking people living in northern Siberia, being shared by ∼90% of Sakha males^72,73^. In our dataset, N1a-M178 Y-chromosomes were found in three Jalayirs and one Shapirashti individuals. By contrast, a closely related clade, N1a2b-P43, characterizes Uralic speakers and other populations in northwestern Siberia and Scandinavia^71,73^. N1a2 also occurs at low frequencies in Central Eurasia, Siberia, Korea, and Japan^71^. The earliest example of N1a2 was found in a Neolithic site near Lake Baikal^74^. Interestingly, one sample in our dataset belonged to N1a2b, having the derived allele of N-Y3199 along with the upstream CTS7235 SNP and other markers.

### Q1-L472

The haplogroup Q1-L472 occurs at high frequencies in Siberia^69^ and its subbranch, Q1b-M3, is the most common paternal lineage found in Native Americans^62^. Q1b-L54, that is ancestral to Q1b-M3, has provided evidence for the prehistoric genetic link between the Altay Mountains and the Americas^69^. In our dataset, Q1-L472 Y-chromosomes were found in only two individuals, one Jalayir (Q1b1a-L54) and one Shapirashti (Q1a1a-F745).

### J-M304

Haplogroup J-M304 likely originated in the Fertile Crescent, with its subbranches J1a and J2a being typical paternal lineages of Arabic and Jewish populations in the Middle East^75^. Thus, their occurrence, at relatively low frequencies, in Kazaks and other populations in Central Eurasia might be associated with the spread of Islam from the Middle East^76^. However, haplogroup J lineages have been discovered in prehistoric burials in Mongolia^77^; therefore, they must have been introduced into Central Eurasia much earlier than the spread of Islam in the 6^th^-7^th^ centuries CE. In our dataset, we observed five J-M304 Y-chromosomes, including J1a2b-CTS15 in an Alban and a Shapirashti, and J2a1a1-L25 in three Jalayirs.

### Other haplogroups

The frequency of I-M170 is very low in Kazaks and other groups in Central Asia^65,78,79^. The highest frequencies of I-M170 occur in Europe, where it likely arose^65^. The Jetisuu Kazaks had just two I-M170 Y-chromosomes, one in each of the Shapirashti and Suan clans.

G2a1a-Z6553 (i.e., G2-P16), which occurs mainly in the Caucasus^80^, was found in two Jetisuu individuals, one in a Shapirashti and another in a Jalayir. Its frequency is generally low in other Kazak and Central Asian populations. However, a closely related lineage, G1-M342, occurs at a considerably high frequency in the Arghin clan of Kazaks^14^, while otherwise being virtually absent in other Kazak populations.

Haplogroup T is a rare lineage in Central Asia^79^, with its possible origin being the Near East^81^. The highest frequencies of T-P77 occur in Kurdish populations and Iraqi Jews^81^. In our sample set, T-P77 (i.e., T1a1a1) occurred in a single Alban individual.

### West Eurasian admixture gradient in Kazaks

As expected, Jetisuu Kazaks were positioned at the center of the PCA plot representing the genetic diversity of human populations in Eurasia (Figure 5). PC1 explained most of the genetic variation in Jetisuu individuals, showing an east-to-west gradient that implied variation in West Eurasian admixture components. Tuba individuals were separated from Kazaks along PC2 despite having comparable positions on PC1, primarily due to their higher Siberian ancestry levels (Figure 6C). The Kirghiz showed the strongest genetic affinity with Kazaks and were positioned eastward along the east-west genetic gradient formed by Kazak and Kirghiz individuals (Figure 5). The PCA analysis did not reveal any pronounced genetic structure in Kazak populations, including the Jetisuu clans and other Kazaks in our dataset. They clearly separated from all other ethnic groups and formed a single cluster in the PC space (Figure 5).

**Figure 5.**
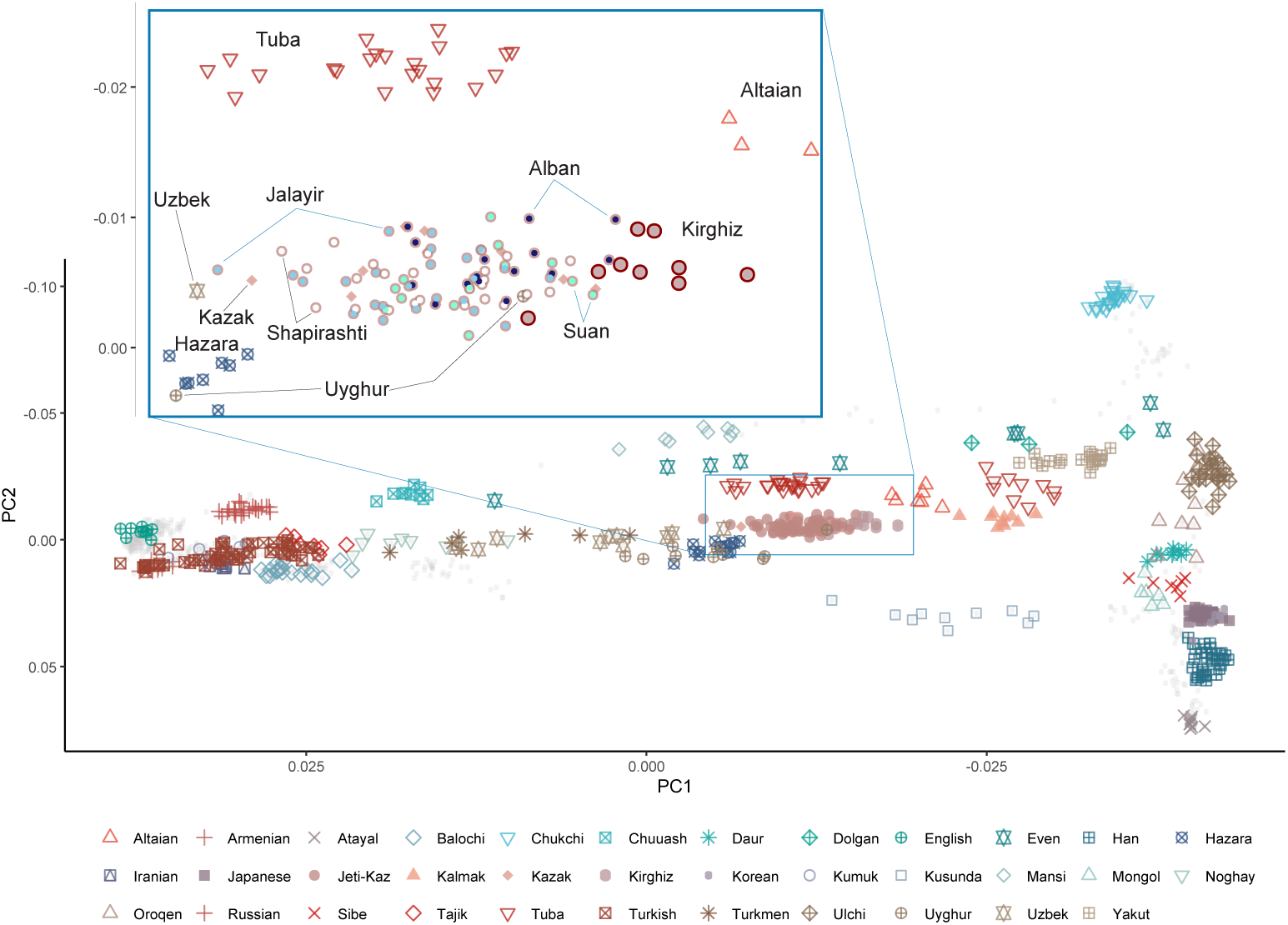
PCA analysis of Jetisuu Kazaks in comparison with the populations across Eurasia. Note: The inset displays Kazaks and other closely related populations, revealing that Kazaks do not exhibit pronounced population structure.

**Figure 6.**
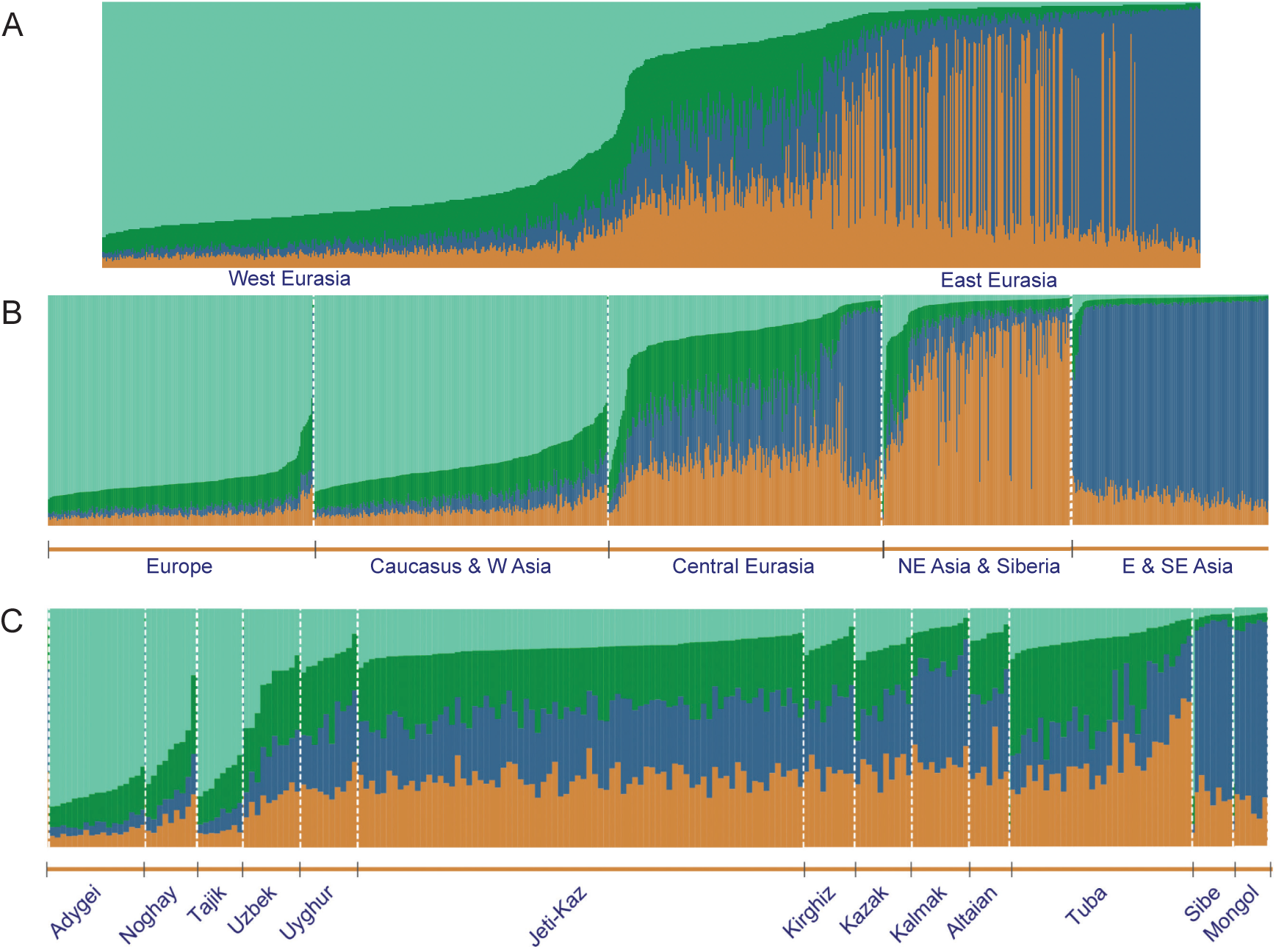
Population admixture in Central Eurasia in the context of the Eurasian genomic landscape. **A.** Individuals across Eurasia, from Japan to England; **B.** individuals grouped by geographic area; **C.** individuals from selected ethnic groups by longitude and/or the West Eurasian genetic component level. **Clusters:** West Eurasian (light green), Southeast Asian (dark blue), Ancient North Eurasian (ANE, dark green), and Ancient North Asian (ANA, orange). **Jeti-Kaz:** Jetisuu Kazaks.

During the historic time periods, three major groups contributed to the formation of Central Asians. These included the Indo-Iranian speaking Sogdians and the Altaic speaking Turks and Mongols, with the latter two being nomadic pastoralists^1,8,82^. The well-known Bronze Age Bactria-Margiana Archaeological Complex (BMAC) culture of Central Asia derived from local Copper Age populations that ultimately trace their ancestry to Iran^66^. The Iron Age Scythians and others prehistoric populations may also have contributed to the genomic profiles of human populations in Central Eurasia^83^. The overall genetic features of Turkic speakers in Central Eurasia can be modeled as various degrees of admixture between Persian speaking Sogdians (or modern Tajiks) and Turko-Mongolian speakers.

### Admixture analysis reveals hiatus in the genetic cline of Central Eurasia

We performed admixture analysis on 1,013 samples using the neural-admixture program^44^. As shown in Figure 6, we could model Central Eurasia as having four ancestral sources (K=4), including Ancient Northeast Asian^84^ (ANA, orange), Ancient North Eurasian^86^ (ANE, dark green), West Eurasian (light green), and Southeast Asian (dark blue). The West Eurasian component (light green) most likely represents ancestry derived from Early European Farmers (EEF), which contributed the majority of the genomic heritage of modern Europeans^42^. However, the analysis at this K-level did not resolve the Western Hunter-Gatherers (WHG) ancestry, that is the second largest genomic component of Europeans^42^; instead, WHG variation was likely subsumed within the broad West Eurasian cluster. We also observed minor traces of ANA and Southeast Asia related sources in Europe, likely reflecting deep Eurasian drift. The ANE ancestry corresponds to the third source of European genomes based on the tripartite-ancestry model^42^, and ANE is represented by the dark green component in the admixture plot (Figure 6).

These ancestral sources for Kazaks and other populations in Central Eurasia are similar to those identified for Kazaks in a previous study^85^. In the current analysis, East Asian groups differed from one another depending on the proportion of Siberian and Southeast Asian ancestry components (Figure 5 & 6), whereas Lei and colleagues^85^ did not differentiate northern and southern East Asians, but identified an additional South Asian ancestral source, which did not appear in our analysis.

The West Eurasian component increases from East to West, as expected. However, this pattern does not follow a gradual gradient along longitudes. Instead, it increases sharply in the Caucasus, with a corresponding decrease in the Southeast Asian, ANE, and ANA components (Figure 6). For instance, the genetic makeup of the Adygei from Caucasus and the Kazaks just east of the Caspian Sea is considerably different (Figure 6C).

These findings suggest that the Caspian Sea and the Caucasus Mountains acted as geographic barriers for gene flow from the eastern steppes. A similar pattern was also observed in southern Central Asia, where Tajiks had an elevated level of West Eurasian ancestry comparable to what was seen in the Caucasus (Figure 6C). Here again, the high slopes of the Tängirtau Mountains served as a geographic barrier for gene flow. Noghays in the Caucasus, who, like Kazaks, derive their ancestry from the Golden Horde^1^, had much higher level of West Eurasian genetic ancestry due to successive phases of admixture with Caucasus populations over the past few centuries (Figure 6C). Similarly, a higher level of West Eurasian ancestry characterizes the Uzbek individuals, who exhibit varying admixture profiles (Figures 5 & 6C).

Together with Tajiks, they form a genetic cline (Figure 6C). Interestingly, this transition zone appears to mark the genomic boundary between Indo-European speakers and Altaic speakers, with highly admixed Turkic speaking populations (e.g., Uzbeks and Noghays in Figure 6C) situated at this interface. Notably, some Turkic speakers (e.g., Anatolian Turks) possess predominantly West Eurasian ancestry similar to neighboring Indo-European speakers (Figure 5).

### Decreased levels of inbreeding in the nomads of Central Eurasia

We analyzed Jetisuu Kazaks (N=78) together with over 900 Eurasian individuals, for the evaluation of Runs of Homozygosity (ROH). ROH indicates the extent of sequence identity between two homologous chromosomes, which are either of maternal or paternal origin. ROH provides us with information about demography, culture, and health of a population^87^.

Individuals from small populations usually exhibit higher levels of ROH due to its members being genetically related to each other. However, members of a relatively large population that recently recovered from a population bottleneck may also show elevated levels of ROH^87^. For example, a Kusunda individual from Nepal had about 80 ROH segments that totaled to more than 250 Mb (Figure S1). Such a large number of shorter ROH segments is indicative of small population size and a genetic bottleneck^87^.

We analyzed the ROH levels of human populations in Central Eurasia, with special reference to the traditional subsistence patterns, e.g., sedentary vs nomadic (Figure 7). We observed pronounced differences in the ROH levels of various ethnic groups, which can be largely categorized into three groups: (1) High ROH group, (2) Medium ROH group, and (3) Low ROH group.

**Figure 7.**
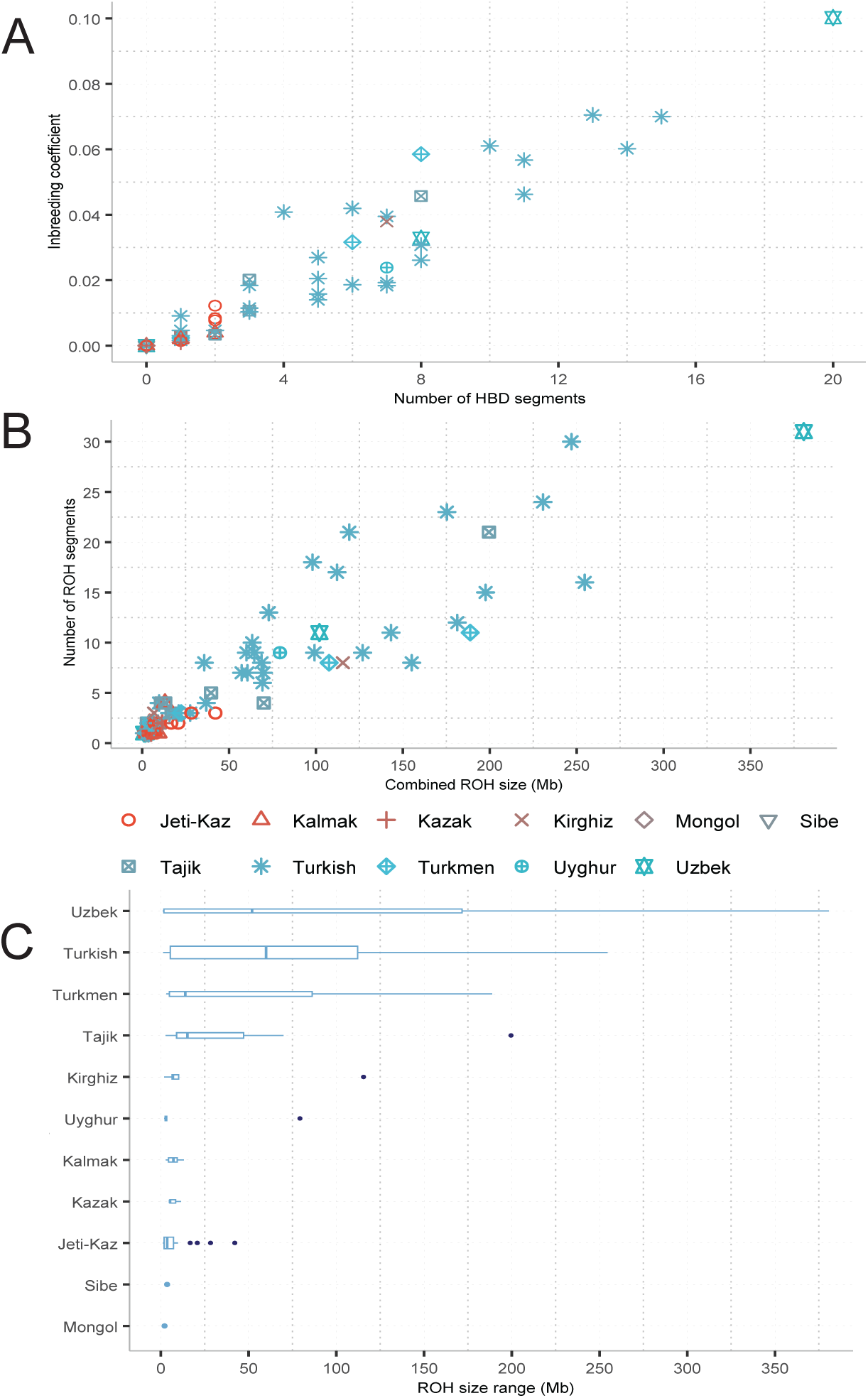
Runs of homozygosity and inbreeding coefficient in Central Eurasian populations. **A.** Inbreeding coefficient obtained from HBD (Homozygosity by Descent) segments with a size of 7 cM or longer; **B.** Contrasting ROH segments by size and count; **C.** Boxplot for all ROH segments; Box width indicates relative sample size, the box length represents quartile values, and the vertical bar shows the median value; the dark blue dots represent outliers with very long ROH. The light blue dots indicate ROH segments found in a single individual of Sibe and Mongol, respectively. In the Boxplot, the populations were ordered based on the mean value of their ROH sizes.

The high ROH group included Turks, Turkmens, Tajiks and Uzbeks, with the latter having the highest mean ROH (Figure 7C). As shown in Figure 7B, an Uzbek individual had about 30 segments, with the combined ROH size being about 375 Mb, encompassing approximately 13.2% of the human genome (based on the genome size of ∼2881 Mb for GRCh37/hg19) and surpassing the cutoff for consanguinity specified by the American College of Medical Genetics and Genomics (ACMG) guidelines^88^. Consequently, the Uzbek individual had an inbreeding coefficient of approximately 0.1, which can result from double first cousin marriages, uncle-niece unions, or first cousin marriages in a community with a complex, multiple generation consanguineous loop^87^.

Consanguinity seems to be common in Uzbeks. A study reported that 33.2% of blind and severely visually impaired Uzbek children were born to consanguineous couples^89^. The Turkish individuals exhibited the second highest levels of ROH (Figure 7C) that was expected given the high rates of consanguineous marriages in Turkey^26^. We have included Turkish individuals in the analysis, since the Turkish people have cultural and historical connections with Central Eurasia.

The moderate ROH group included a number of Turkic speaking groups from Siberia. The Sakha (Yakut) and Tuba had moderate levels of ROH, with the combined length of ROHs was less than 100 Mb, featured by multiple number of shorter ROH segments (Figure S3). This outcome was expected, given the fact that native groups in Siberia usually have small population sizes and previously experienced population bottlenecks. By contrast, the Tajik, Turkish, Turkmen, and Uzbek individuals had fewer but very long segments with a combined length of 150 Mb or longer, suggestive of consanguinity and inbreeding (Figure 7, S3, and S4).

Contrasting sharply with the ROH found in sedentary agriculturalists in Central Eurasia, such as Turkmens, Tajiks, and Uzbeks in Figure 7, the low ROH group included the pastoral nomads—Kalmaks, Kazaks, and Mongols---and the Tungusic speaking Sibe (Figure 7). The Kalmaks and Mongols are Mongolic speakers and practice Buddhism, but share pastoral nomadic culture with Kazaks and Kirghiz. The four also had similar traditional belief systems prior to accepting Buddhism or Islam. As previously mentioned, the pastoral nomadic groups have low rates of consanguineous marriages (Figure 2H), which is reflected in their genomes as low levels of ROH (Figure 7 and Figure S4).

In our dataset, the nomadic Kirghiz had a higher mean ROH than that of the sedentary Uyghurs (Figure 7). This might be due to not having sufficiently representative samples from these populations. Despite this possibility, a considerably high level of consanguinity was reported for some Kirghiz communities in Kirghizstan (1.4 – 4.0% for the older generation, and 5.8 – 12% for the younger), even though exogamy is still predominant among the Kirghiz in general^90^. The increased rate of consanguinity could reflect the shift from a nomadic to a sedentary lifestyle, as well as the increased Islamization of the younger generation.

We categorized the populations into two cultural groups—pastoral nomads and sedentary agriculturalists. The nomads included Jetisuu Kazaks, Kazaks (*i.e., Kazaks from other regions*), Kalmaks, Kirghiz, and Mongols, whereas the agriculturalists were composed of Turkmen, Tajik, Uzbek, and Uyghur individuals. We did not include Turks and the Sibe in the two cultural groups because the former come from a different geographic region while the latter cannot be easily classified as either a nomadic or a sedentary group. Originally from Manchuria, the Sibe were stationed at their current location as a western frontier garrison by the Manchu Empire in the 18^th^ century^91^.

Using a Mann-Whitney U (MWU) Test and Fisher’s Exact Test, we evaluated ROH differences between the two cultural groups. For the tests, we only included individuals having at least one ROH segment, and this reduced our sample set to 49 nomads and 23 agriculturalists for comparison. The total sizes of the ROH segments showed statistically significant differences when all individuals with at least one ROH segment were included (MWU W=749 with p-value=0.0247). Again, the difference was significant when the proportion of individuals with long ROH segments (34.8Mb or longer) were compared (Fisher’s Exact Test p-value=0.001152). The test results indicate that, in contrast with the sedentary agriculturalists, pastoral nomads in Central Eurasia have significantly lower levels of ROH.

Kazaks are historically nomadic and intentionally avoided marriage between close relatives, in particular, paternally related individuals. This exogamic practice encouraged marriage between two individuals who were at least seven generations apart as reckoned through the paternal lineage. Nowadays, exogamy is still practiced in Kazak society, despite no written law that explicitly stipulates it. This marriage pattern is part of Kazak culture, and closely linked with the clan-based social structure and genealogy tradition. The Soviet regime had a failed attempt to dismantle the Kazak social structure^92^.

Despite these observations, it takes just a few generations for nomadic peoples to lose their traditional values after adopting a sedentary lifestyle^93^ and intermarrying with locals. Consistent with this trend, we observed higher West Eurasian admixture components in the sedentary Turkic speakers (Figure 5 and Figure 6C). Even though nomadic Seljuks and Özbeks were responsible for the formation of modern Turks and Uzbeks, respectively, Muslim cultural practices have largely replaced the traditional nomadic ones in these societies. Thus, paternal parallel first-cousin unions and other types of consanguineous marriages are now allowed^26^, mirroring marital practices common in the Middle East and North Africa, where Islam prevails^94,95^. The same applies to the other two Turkic speaking groups—Turkmens and Uyghurs—even though the Turkmens practiced nomadic or semi-nomadic pastoralism well into the 20^th^ century^96^.

Our findings are in alignment with the low levels of ROH and inbreeding coefficient previously found in Kazaks from Uzbekistan^97^. The ROH levels of Uzbeks reported by the same study were low as well, indicating that consanguineous marriages are not evenly distributed in Uzbekistan, occurring more in some regions or communities. Furthermore, Marchi and colleagues found high levels of ROH in the group of Turko-Mongolian populations that included Kazaks, Uzbeks, and a few other ethnic groups from Siberia. Higher levels of ROH were expected for the Turko-Mongolian group in this study, since most of them (7 out of 12) were sampled from southern Siberia, where many populations experienced population bottleneck, with small population size and relatively low genetic diversity, due to previous population bottlenecks. Even though marriage unions were formed through geographic exogamy, many of the couples were second cousins (e.g., 40% in Telengits, 69% in Shors, and 76% in Tuba)^97^. When all Turko-Mongolian and Indo-Iranian populations were compared, Tajiks from Dughoba, Tajikstan had the highest mean value of inbreeding, with highest number of long ROH segments^97^, implying consanguinity, as the case of Uzbek individuals in our dataset.

### Kinship among the Jetisuu individuals

We estimated the kinship coefficient among the Jetisuu Kazaks and other individuals of various ethnic backgrounds in our dataset, based on shared IBD segments retrieved via the IBIS program^53^. We then reconstructed a Neighbor-Joining tree for the individuals with up to a 7^th^ degree of relatedness (Figure S5). As expected under an exogamy model, most pairs or trios who clustered as kin had different clan or Jüz identities, with a few even coming from other Turkic speaking nomadic groups, such as Kirghiz and Noghay, reflecting wider exogamous network.

Like Kazaks, Kirghiz and Noghay speak languages from the Kipchak branch of Turkic and share historical connections. The close kin individual pairs who belonged to the same clan constituted about one third of these pairs. In addition, most of the kin pairs had 7^th^ degree relatedness (∼90%), with only few having closer kinship (4–6^th^ degree, Table S4). Several Jetisuu individuals did not have up to the 7^th^ degree of relatedness with any individuals in the dataset (Table S4).

## Discussion

The evolution of human genes has been influenced profoundly by technological and cultural factors, with cultural transmission being an important human social behavior^98^. As a pastoral nomadic group in Central Eurasia, Kazaks traditionally orally transmitted the knowledge of social structure and genealogy from generation to generation.

Our study addresses two fundamental questions about nomadic social organization in Central Eurasia. First, we asked whether the paternal lineage-based genealogy of nomads matches their Y-chromosome phylogenetic relationships. Our results demonstrate substantial concordance between genetic and genealogical data at the clan level in the Jetisuu Kazaks, with most individuals within a clan sharing one dominant paternal lineage (C2a1a3-F1918) that aligns with their claimed ancestry by genealogy. Second, we examined the genomic manifestations of social structure and exogamy in the pastoral nomadic populations of Central Eurasia. We found evidence of sex-biased admixture patterns and decreased levels of ROH in Jetisuu Kazaks, consistent with patrilineal social organization and female exogamy. Our findings thus reveal the effects of gene-culture co-evolution, where religion and culture influenced genomic profiles of individuals in a population.

### Reconciling genetic evidence with anthropological theory

These findings contribute to ongoing debates about the authenticity of nomadic genealogies. While anthropologists and historians have argued that clan affiliations reflect fluid political alliances rather than genuine kinship^5,8–10^, our genetic data suggest a more nuanced picture. It was obviously a misconception to believe that, among Kazaks, only aristocrats kept genealogies^9,11^. Most of our research participants in Jetisuu could retrieve the names of seven forefathers either from memory, by checking a genealogy book, or by consulting a relative. At the individual clan level, we observe strong correspondence between genetic and genealogical identity, supporting the biological reality of paternal lineages. However, we also detect secondary lineages within these clans, such as haplogroup R1a1a among Suans, which may represent more recent admixture events. This pattern further suggests that, while the core genetic structure of clans has persisted over time, the kinship system also accommodated political and social integration of non-relatives.

### Gene-culture co-evolution in nomadic societies

The anthropologist William Durham has proposed incest taboos as an example of gene-culture co-evolution in human populations that have such taboos^99^. He asserted that incest taboos have been culturally selected due to the fitness costs involved in consanguinity; and thus have enhanced human survival and reproduction^99^. His analysis also indicates that exogamous populations have much stricter incest taboos compared to populations that practice endogamy^99^. Following the same line of reasoning, we believe that our findings demonstrate gene-culture co-evolution in pastoral nomads, where cultural practices of exogamy and patrilineal organization have shaped genetic diversity over many generations. The practice of clan exogamy, reinforced by genealogical knowledge spanning seven generations, functions as a biological mechanism to minimize inbreeding and maintain genetic diversity. Awareness of clan identity and the genealogical knowledge is prerequisite for preserving the tradition of seven-generation exogamy.

The genealogy and exogamic practice reinforce each other. Kazaks use a proverb — “*He who does not know his seven forefathers is an orphan*”— to encourage children to learn genealogy. Similar sayings exist in other nomadic groups, including Kalmaks and Kirghiz. This cultural institution effectively counteracts the Wahlund Effect that would otherwise reduce heterozygosity in isolated groups^100^. The contrast between homogeneous paternal lineages and the previously observed heterogeneous maternal lineages^37^ in Jetisuu Kazaks reflects this sex-biased gene flow pattern, where women move between clans while men maintain their patrilineal affiliations within them.

It is hard to ascertain the exact timing of the origin of exogamy in pastoral nomads in Central Eurasia. However, in Kazaks this marriage custom certainly predates the emergence of Kazaks as a distinct nomadic group in the 15^th^ century CE. We know from historical annals that Temujin (i.e., Genghis Khan) the Borjigin married Börte the Kongirat (Ongirat)^6^. Thereafter, the descendants of Genghis Khan practiced exogamy as well, with the Kongirat being the main source of women^101^. In addition, ancient DNA studies indicate the existence of exogamy in the medieval pastoral nomads. For example, the medieval Avars from the Carpathian Basin exhibited short runs of homozygosity (ROH), suggesting that they had intentionally avoided consanguineous marriages^102^. The marriage practice of avoiding close paternal relatives may possibly trace its origins back to prehistoric Altaic speakers in Siberia or Northeast Asia, who organized themselves as patrilineal descent groups. Future studies may reveal the exact timeline of the development of exogamy in the pastoral nomads.

### Evolutionary and Social Implications of Exogamy

The persistence of exogamous marriage practices in nomadic societies offers both biological and social advantages. Genetically, it prevents the accumulation of deleterious recessive alleles and maintains adaptive potential through continued gene flow. Exogamy promotes the genetic benefits of sexual reproduction by augmenting within-individual as well inter-individual genetic heterozygosity, while inbreeding counteracts these benefits by increasing homozygosity^99^. Socially, inter-clan marriages create extensive networks that enhance cooperation and political stability across clan confederations. This dual benefit could explain why such systems evolved and persisted despite the organizational complexity required to maintain complex genealogical records across multiple generations. The seven-generation exogamy rule of pastoral nomads represents a balance between preventing close inbreeding while maintaining clan identity and social cohesion.

### Complexity of nomadic social organization

While our data support a genetic foundation for clan structure, several observations highlight the complexity of nomadic social organization. For one thing, the genetic basis becomes less clear at higher organizational level, i.e., the *Jüz*. As an example, the Middle Jüz encompasses six clans (Figure 1) with diverse paternal origins (G1-M342 in Arghins^14^, O-M134 in Naymans^103^, C2b-M407 in Kongirats^56^, and C2a-F1918 in Kereys^13^), suggesting that larger military or administrative unions may indeed represent more fluid political and social alliances. Through this higher-level social structure—Jüz—six separate patrilineal descent groups achieved an effective union, as seen in the formation of League of Iroquois by six Native American nations^3^. The Iroquois nations practiced exogamy, whereby the husband and wife belonged to different clans, although the clan identity of the children followed the maternal lineage. As a result, the father and son each belonged to distinct clans^3^.

Additionally, the presence of minor haplogroups within Kazak clans indicates that the system accommodated adoption, political integration, or recent admixture events. Such patterns suggest that nomadic genealogies reflect both biological kinship and social construction and thus cannot be characterized as purely genetic or completely fictitious.

### Contrast between pastoral nomads and sedentary agriculturalists

Our analysis reveals a sharp contrast between the two cultural groups regarding marriage patterns and ROH levels in the genome. Contrary to our results, a previous study found higher overall levels of ROH in Turko-Mongolians than in Indo-Iranians (Tajiks)^97^. However, this observed difference is likely attributable to an abundance of shorter ROH segments in the former group. These short segments accumulated due to historical population bottlenecks and small effective population sizes, as most populations in that study’s Turko-Mongolian group originated in Siberia (as the characteristics of the Siberians in Figure S4). Furthermore, grouping population solely by linguistic affiliation can create analytic problems. For instance, the Turkic speaking Uzbeks are culturally as well as genetically closer to the Indo-Iranian speaking Tajiks than they are to the Tuba and other Turkic speaking groups in Siberia (Figure 6C).

The sedentary agriculturalists follow Islamic institutions and frequently practice consanguineous marriages. While often associated with Islam, the consanguinity might have existed in Central Asia and neighboring regions prior to the religion’s spread, with Islam subsequently reinforcing the custom. In this regard, an Iron Age individual from Otirar (southern Kazakstan) had long ROH segments totaling 300 Mb, a clear signature of consanguinity and inbreeding^104^. Similarly, an Iron Age individual from northwest Pakistan exhibited long ROH resulting from a consanguineous marriage^105^. Despite these sporadic ancient occurrences in the Middle East, West Asia, and Central Asia, though, modern ROH levels are much higher than those of ancient populations in these regions^105^.

Notwithstanding the apparent genetic and health risks, consanguineous marriages persist, likely due to their perceived social benefits. These include: (1) the maintenance of social structure and house property; (2) simplified marriage arrangements with reduced financial burden and greater marital stability; (3) uniting the individuals with the same descent; and (4) keeping tradition and transmitting cultural values, etc^95^.

### Future Directions

Our findings demonstrate that the social structure of Central Eurasian nomads represent sophisticated examples of gene-culture co-evolution, where cultural institutions have systematically influenced genetic patterns over centuries. Future research using high-resolution Y-chromosome sequencing may potentially provide more precise estimates of divergence times and admixture patterns within and between clan lineages. In addition, the ROH characteristics of various groups in Central Eurasia could be further consolidated by surveying larger samples with higher representative and statistical power. The integration of autosomal data would further illuminate the complex admixture history of these populations. More broadly, this study illustrates how population genetic approaches can illuminate the biological reality underlying cultural institutions, providing new perspectives on long-standing anthropological debates about the nature of kinship and social organization in pastoral nomadic societies.

## Supplemental Information

**Supplemental material:** Contains Figures S1 through S5.

**Table S1.** Jetisuu samples and their Y-chromosome haplogroups.

**Table S2.** Information on the ROH segments found in 1013 individuals across Eurasia, including Jetisuu Kazaks.

**Table S3.** Information on the inbreeding levels of 71 populations across Eurasia, including Jetisuu Kazaks.

**Table S4.** Kinship Coefficients of the Jetisuu kin pairs with up to 7th degree relatedness.

## Declaration of Interests

The authors declare no competing interests.

## Acknowledgments

We sincerely thank the individuals who participated in this study by providing their samples. Our appreciation extends to the Center for Alash History in Almati city for assisting the fieldwork. We are also grateful to the local government authorities of Almati and Jetisuu Oblast, as well as the county and village administrations, for their support in facilitating the field activities. Additionally, we acknowledge the local hospitals in Almati and Jetisuu Oblast for their cooperation in supplying nurses to collect blood samples. Fieldwork and DNA preparation lab work of the project were supported by a Targeted Funding Program for 2014-2017 from the Ministry of Education and Science of the Republic of Kazakstan, which was awarded to the Center for Life Science at NU. The GenoChip microarray analysis was conducted as part of The Genographic Project, supported by the National Geographic Society,

USA. Finaly, we appreciate the historian Peter Golden for reviewing the manuscript and providing valuable comments.

## Web Resources

NA

## Data and code availability

GenoChip microarray data of 90 individuals will be available at GEO upon publication.

## Notes

### Competing Interest Statement

The authors have declared no competing interest.

