## Supplemental Data 1 for "Genetic Basis of Social Structure in the Pastoral Nomads of Central Eurasia"

### Supplemental Material

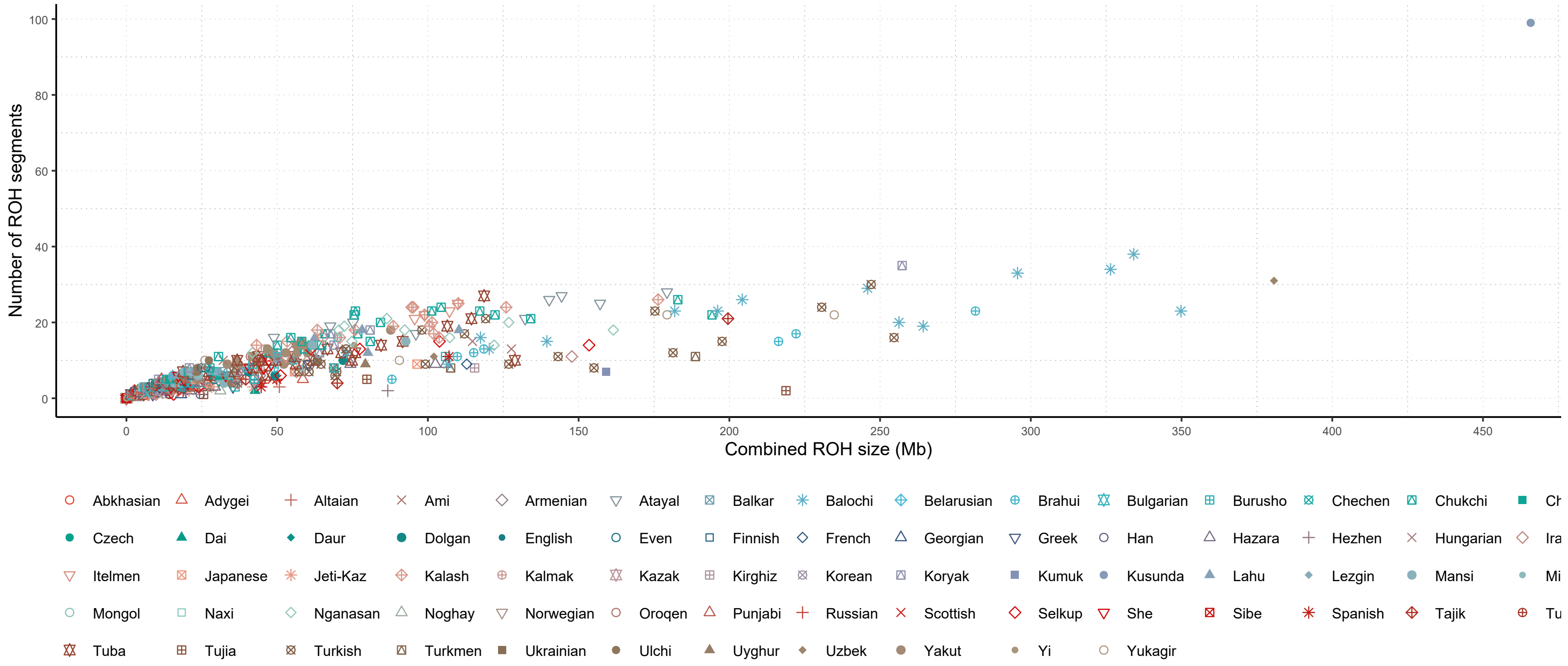


Figure S1. Runs of homozygosity (ROH) levels observed in Eurasian populations.

A total of 1013 individuals from 71 Eurasian populations was analyzed for ROH via PLINK v1.9[1]. Each data point in the scatter plot indicates an individual that belongs to a population in the list given under the plot. Y-axis indicates the total number of ROH segments found in the genome of an individual. X-axis shows the combined size of all ROH segments in an individual. The scatter plots were made via ggplot2[5] in R[4].

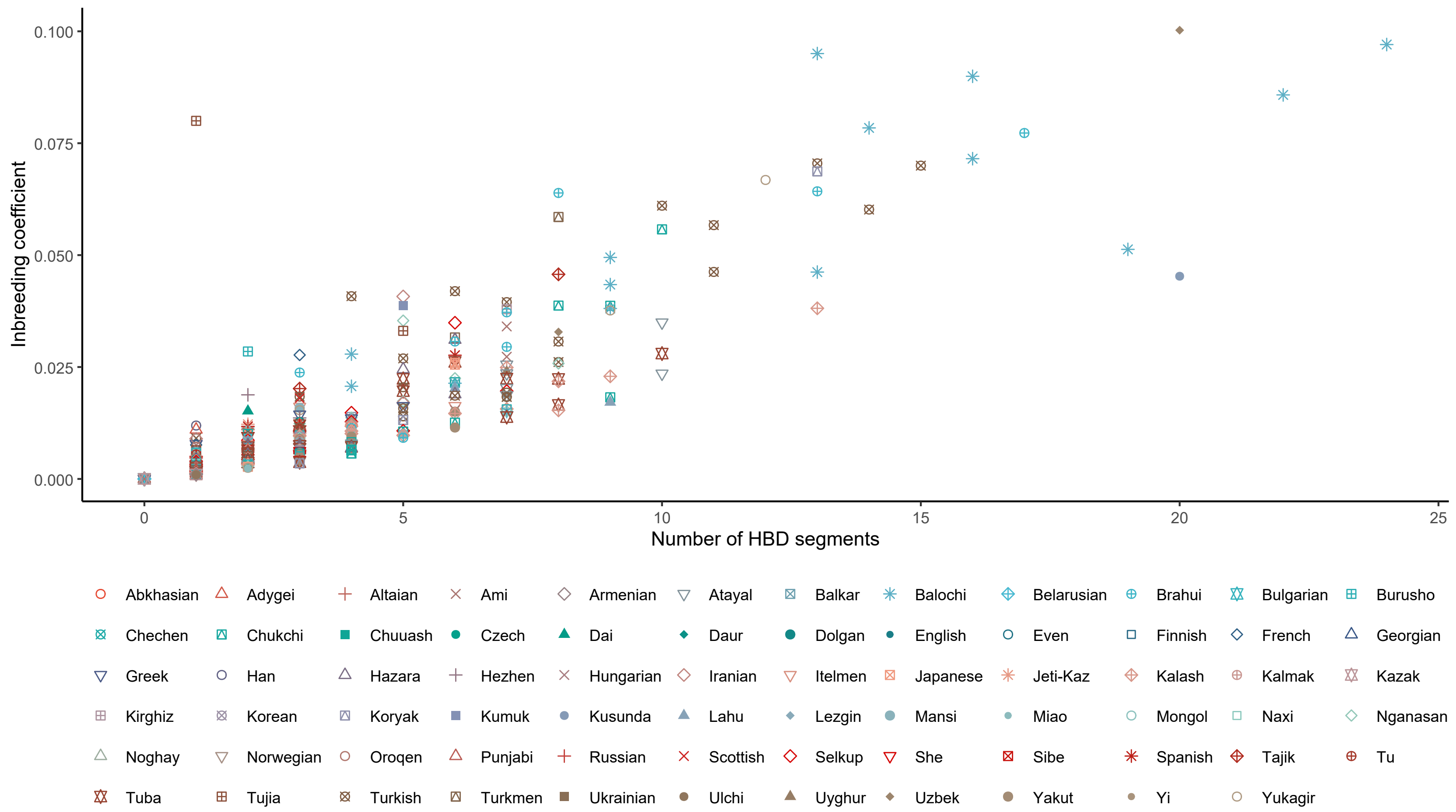


Figure S2. Inbreeding levels observed in Eurasian populations.

A total of 1013 individuals from 71 Eurasian populations was analyzed via the IBIS program[2]. Each data point in the scatter plot indicates an individual that belongs to a population in the list given under the plot. IBIS detects longer ROH segments compared to PLINK, therefore, total number of segments of many individuals shown in this figure are less than those of Figure S1. This is particularly conspicuous for the Kusunda individual with the longest ROH in Figure S1. Y-axis indicate inbreeding coefficient calculated based on homozygous sites found in the genome of an individual. X-axis shows the number of HBD (Homozygosity by Descent, i.e., Runs of Homozygosity) segments in an individual. The scatter plots were made via ggplot2[5] in R[4].

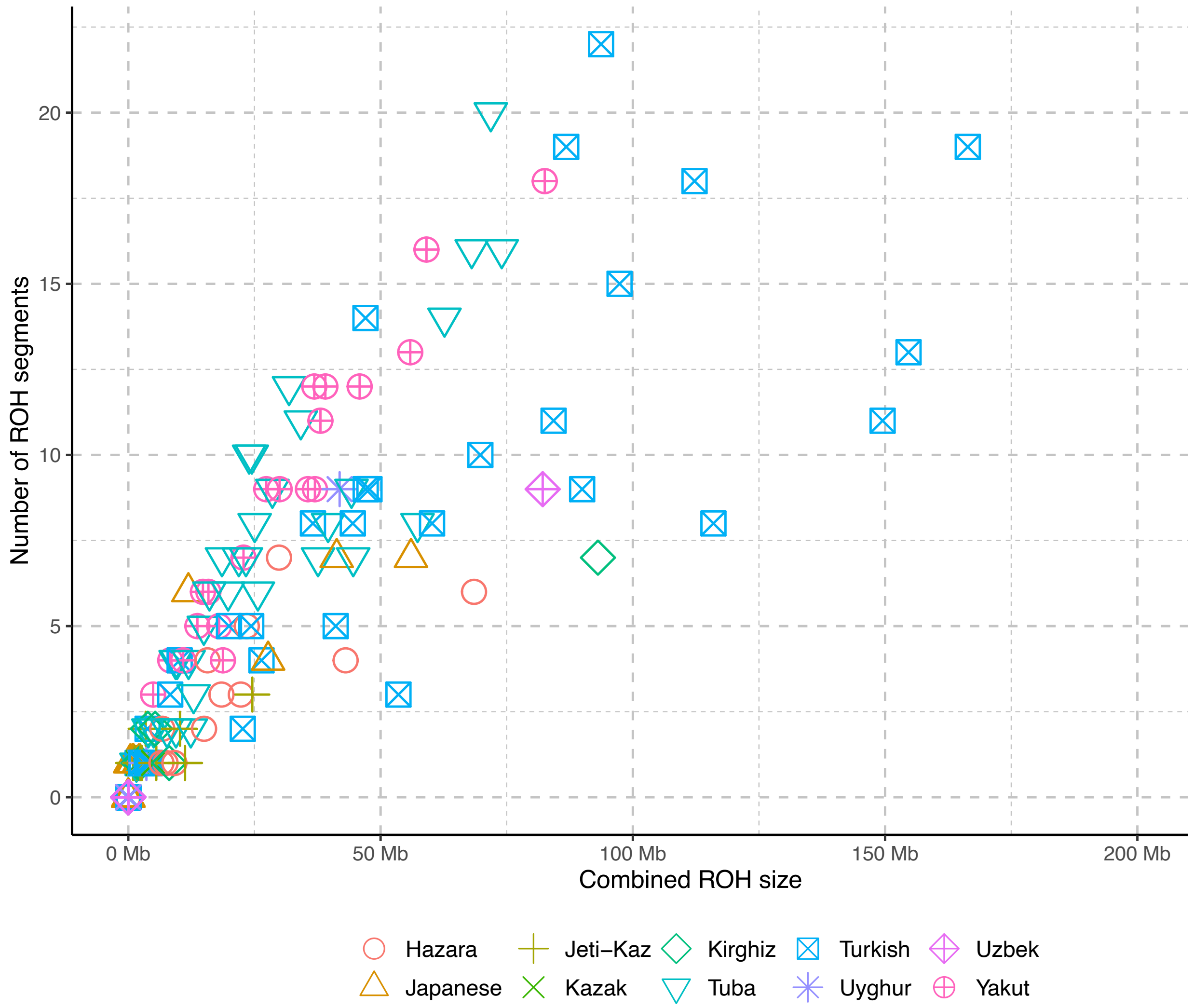


Figure S3. Total Runs of Homozygosity (ROH) segments found in a selected set of populations.

Y-axis indicates total number of ROH segments while X-axis shows their combined sizes in Mb.The scatter plot was made via ggplot2[5] in R[4].

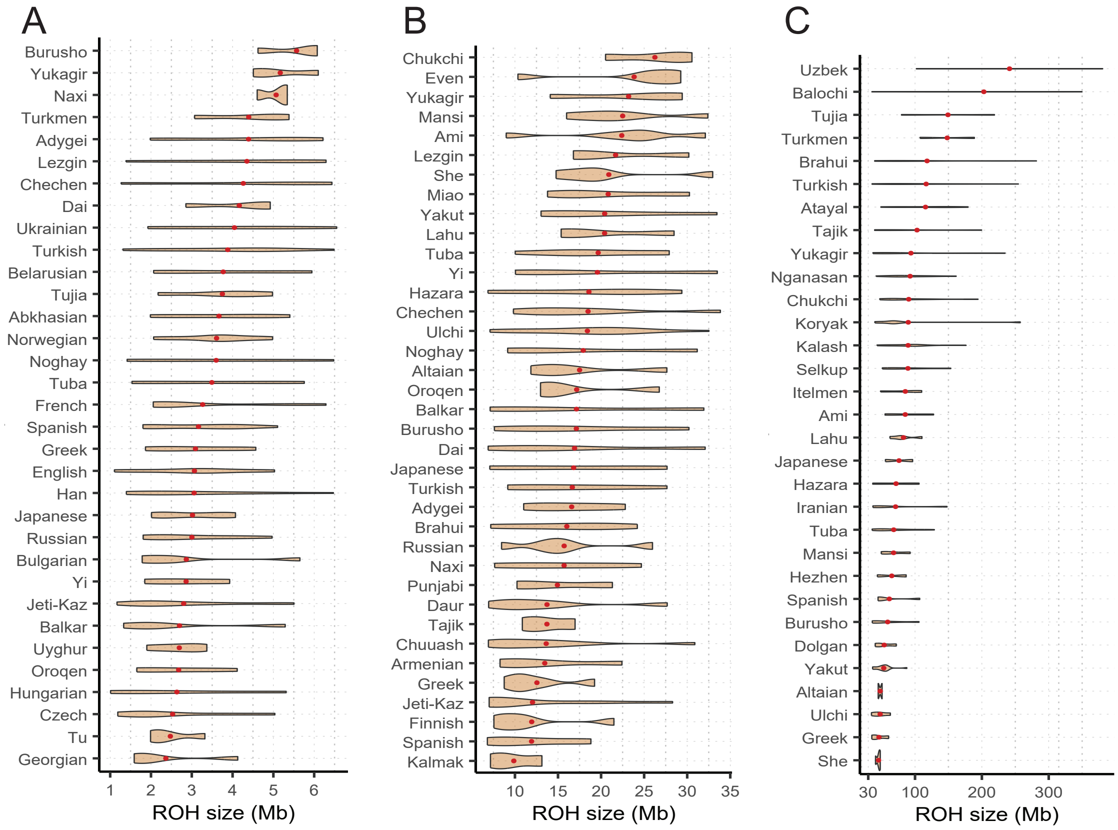


Figure S4. Total Runs of Homozygosity (ROH) levels observed in Eurasian populations.

A. Low ROH (1–6.5Mb, 33 populations with 3 or more individuals were included for the visibility of density); B. Medium ROH (6.7–33.8Mb, 37 populations with 3 or more individuals were included for the visibility of density); C. Long ROH (34.8–465.8Mb, 31 populations with 2 or more individuals were included). The populations were ordered by mean ROH sizes. A total of 1013 individuals from 71 Eurasian populations was analyzed. No ROH was detected in 281 individuals, whereas the ROH segments detected in 732 individuals (representing all 71 populations) were clustered into three categories via the Mclust method[3] in R[4]. The violin plots were made via ggplot2[5] in R[4].

Figure S5. IBD based kinship among Jetisuu and other individuals

Notes: The kinship tree was constructed among individuals with up to the 7th degree of relatedness. Green clusters indicate the relatedness within the same clan, given shared paternal ancestry; Brown clusters show kinship between Jüz; Pink clusters indicate kinship network between ethnic groups; All other clusters without color show kinship among the Great Jüz clans. The figure was created via the R package ggtree[6].

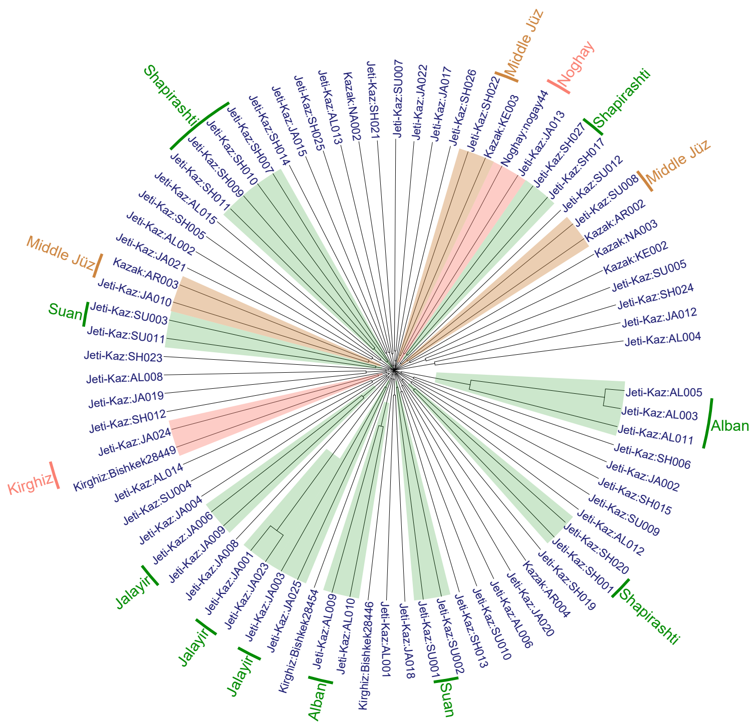
